# Diverse Arctic lake sediment microbiota shape methane emission temperature sensitivity

**DOI:** 10.1101/2020.02.08.934661

**Authors:** Joanne B. Emerson, Ruth K. Varner, Martin Wik, Donovan H. Parks, Rebecca B. Neumann, Joel E. Johnson, Caitlin M. Singleton, Ben J. Woodcroft, Rodney Tollerson, Akosua Owusu-Dommey, Morgan Binder, Nancy L. Freitas, Patrick M. Crill, Scott R. Saleska, Gene W. Tyson, Virginia I. Rich

## Abstract

Northern post-glacial lakes are a significant and increasing source of atmospheric carbon (C), largely through ebullition (bubbling) of microbially-produced methane (CH_4_) from the sediments^1^. Ebullitive CH_4_ flux correlates strongly with temperature, suggesting that solar radiation is the primary driver of these CH_4_ emissions^2^. However, here we show that the slope of the temperature-CH_4_ flux relationship differs spatially, both within and among lakes.

Hypothesizing that differences in microbiota could explain this heterogeneity, we compared site-specific CH_4_ emissions with underlying sediment microbial (metagenomic and amplicon), isotopic, and geochemical data across two post-glacial lakes in Northern Sweden. The temperature-associated increase in CH_4_ emissions was greater in lake middles—where methanogens were more abundant—than edges, and sediment microbial communities were distinct between lake edges and middles. Although CH_4_ emissions projections are typically driven by abiotic factors^1^, regression modeling revealed that microbial abundances, including those of CH_4_-cycling microorganisms and syntrophs that generate H_2_ for methanogenesis, can be useful predictors of porewater CH_4_ concentrations. Our results suggest that deeper lake regions, which currently emit less CH_4_ than shallower edges, could add substantially to overall CH_4_ emissions in a warmer Arctic with longer ice-free seasons and that future CH_4_ emission predictions from northern lakes may be improved by accounting for spatial variations in sediment microbiota.

## Main text

At high latitudes, lakes and ponds are recognized as a large and understudied source of methane (CH_4_)^1,3,4^, a radiatively important trace gas. Post-glacial lakes (formed by glaciers and receding ice sheets, leaving mineral-rich sediments) represent the largest lake area at high latitudes^5^. Because of their areal extent, these lakes contribute to approximately two-thirds of the model-predicted natural CH_4_ emissions above 50**°** N latitude^1^. Their geochemistry and emissions are distinct from thermokarst lakes formed by permafrost thaw^6^. With warming, permafrost thaw, and predicted increased precipitation, northern lakes are expected to receive more terrestrially-derived carbon, likely increasing their carbon dioxide (CO_2_) and CH_4_ emissions^7,8^.

Ebullition commonly accounts for > 50%, sometimes > 90% of the CH_4_ flux from post-glacial lakes, with the remainder primarily attributed to diffusion-limited hydrodynamic flux^9,10^. Ebullition moves CH_4_ rapidly from sediments directly to the atmosphere, typically bypassing microbial CH_4_ oxidation in the water column^11^. Incoming short-wave radiation and sediment temperature have been identified as strong predictors of ebullitive CH_4_ emission from sub-arctic post-glacial lakes on an annual basis, with higher temperature increasing emissions during the ice-free season^2,12^. However, the extent and drivers of spatial variability in this temperature response, particularly within lakes, are poorly understood.

To address this knowledge gap, we analyzed CH_4_ emissions over a six-year period and collected underlying sediment cores in July 2012 from the littoral (“edge”) and pelagic (“middle”) locations of two shallow post-glacial lakes, Mellersta Harrsjön and Inre Harrsjön, (Figure S1, Supplementary Table 1). These lakes are part of the Stordalen Mire complex, a hydrologically interconnected, discontinuous permafrost ecosystem encompassing post-glacial lakes and a mosaic palsa/wetland in approximately equal portions^13^. The lakes contribute ∼55% of the total ecosystem CH_4_ loss^2^ and are model sites for studying ebullitive emissions, which were collected at lake surfaces for the six summers from 2009-2014^12,14^ every 1-3 days^9^. Here, we linked site-specific (lake edge vs. middle) CH_4_ emissions to analyses of the microbiota and biogeochemistry in the underlying sediments.

Previous work has shown that annual ebullitive emissions are consistently higher from these lakes’ shallow littoral zones than their deeper pelagic zones^9,15^, as expected, since the shallow sediments experience higher temperatures for longer periods and also receive more substrate input from aquatic vegetation^16^. However, assessing the temperature *sensitivity* of ebullition for the two lake zones in this study revealed a previously unnoticed significant difference, with ∼5-fold higher temperature sensitivity in lake middles relative to edges (Figure 1, Supplementary Table 2). Predicted future emissions from post-glacial subarctic lakes are based on current measurements of temperature responsiveness^1^, which are dominated by ebullitive flux data from shallow lake edges because those locations currently experience a longer period of sufficient warmth for seasonal emissions than lake middles (∼3 months relative to ∼1 month)^2^. If, as suggested here by our spatially resolved emissions data, temperature responsiveness is substantively higher in the deeper sediments, then, as deeper regions warm and remain heated for longer before cooling off in the fall, future lake emissions would be greater than currently predicted. Thus, accurate CH_4_ emission predictions rely on understanding the spatial heterogeneity and underlying causes of this temperature responsiveness.

**Figure 1.**
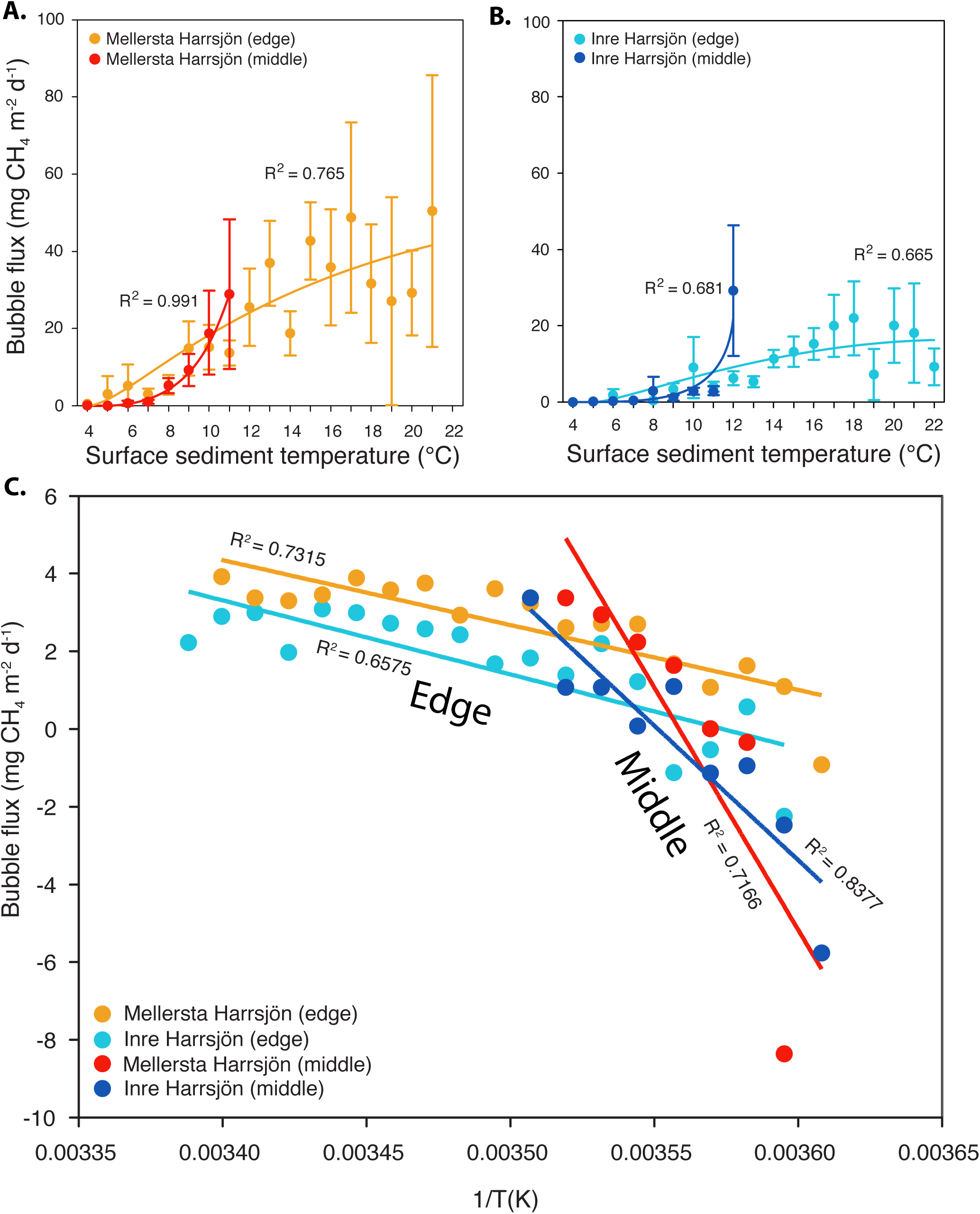
Temperature responsiveness of ebullitive methane flux from two post-glacial lakes. Ebullitive CH_4_ flux as a function of surface sediment temperature (data were binned in 1 °C intervals; see methods) for the edge versus middle regions of: **A.** Lake Mellersta Harrsjön (MH) and **B.** Lake Inre Harrsjön (IH), from June - September 2009 - 2014; MH edge - *n* = 1,609, MH middle - *n* = 810, IH edge - *n* = 2,347, IH middle - *n* = 549. Error bars are 95% confidence intervals, fit lines are 2nd degree polynomials. **C.** Arrhenius plots of the data in A & B; ln (bubble CH_4_ flux) versus the inverse surface sediment temperature in K. Data are color-coded by lake and by edge and middle areas.

Ebullition is controlled by CH_4_ production (which is in turn driven by redox, substrates, temperature, and microbiota), consumption (driven by redox and microbiota)^17-19^, and the physics of bubble formation and escape (determined by sediment texture and overlying hydrostatic pressure, which is largely controlled by atmospheric conditions)^2,15^. Therefore, the edge-to-middle difference in temperature responsiveness of CH_4_ ebullition could be partly due to differences in physicochemical characteristics (*e.g*., sediment texture, pressure, and redox), substrates (*e.g*., organic carbon), and/or microbiota (abundance, composition, and/or activity)^20^. Although differences in sediment texture were observed between the lake edge and middle in Mellersta Harrsjön, these differences were not consistent between lakes (Figure S2, Supplementary Table 3). Our previous work has shown higher and more variable ebullition rates during periods of dropping atmospheric pressure, but there were no differences in edge versus middle locations^9^. In terms of redox, we expect concentrations of terminal electron acceptors to be low, as the likely source would be runoff^21^, and total sulfur and nitrogen did not correlate with ebullition rates by lake or location^15^. In terms of measured substrates, carbon:nitrogen (C:N) ratios and bulk ^13^C_TOC_ (indicative of vegetation composition) did not vary from edges to middles. Total organic carbon (TOC) varied by lake, with similar concentrations observed between lake edge and middle in Mellersta and appreciably higher TOC in middle sediments in Inre Harrsjön. Carbon quality, as assessed by visual comparisons of organic matter composition, revealed coarse, less decomposed detritus gyttja (organic-rich, peat-derived mud) in the edge sediments of both lakes, while middle sediments were characterized by fine-grained, generally more decomposed detritus gyttja^15^. Thus, higher temperature responsiveness occurred where there was lower potential substrate quality, suggesting that substrate differences do not readily explain differences in CH_4_ emission responses to temperature in edge versus middle lake locations, although more detailed substrate analyses could further evaluate this in future.

Next, we sought to characterize differences in microbiota that could contribute to the observed temperature response differences in CH_4_ emissions. We used a 16S rRNA gene amplicon sequencing approach to characterize microbial community composition from the edge and middle cores from each lake (Figure 2A-B, Supplementary Table 4). Although microbial community composition differed most significantly by depth within the sediment (Figure S3, Supplementary Table 5), as is typical for aquatic sediments^22^, significant differences between lake edges and middles (Figure 2C, PERMANOVA *p* = 0.001) suggest that microbiota could contribute to the observed temperature sensitivity in CH_4_ emissions. Indeed, methanogens (defined here as populations from known methanogenic clades^23^, Supplementary Table 4) were significantly more abundant in lake middles than edges (Figure 2D, ANOVA *p* = 0.0001), while total microbial abundances correlated most strongly with depth and did not exhibit edge vs. middle differences (Figure S4, Supplementary Table 6). Aerobic methanotrophs, which are posited to have minimal impact on ebullitive loss due to rapid bubble movement through sediment^11^, were confined to the surface sediment layers as expected (Supplementary Table 7) and did not differ significantly in composition or relative abundance between edges and middles (ANOVA *p* = 0.76). Anaerobic methanotroph abundances differed significantly between lake edges and middles (ANOVA *p* = 0.014, Supplementary Tables 7-8) and were approximately one order of magnitude higher in edge sediments. Although this could suggest that increased anaerobic methane oxidation in the edge sediments could contribute to the observed differences in temperature sensitivity, these anaerobic methanotrophs comprised only 0.1% of the community on average (up to 0.6%, Supplementary Tables 4 and 7), and ebullition is expected to largely bypass methane oxidation.

**Figure 2.**
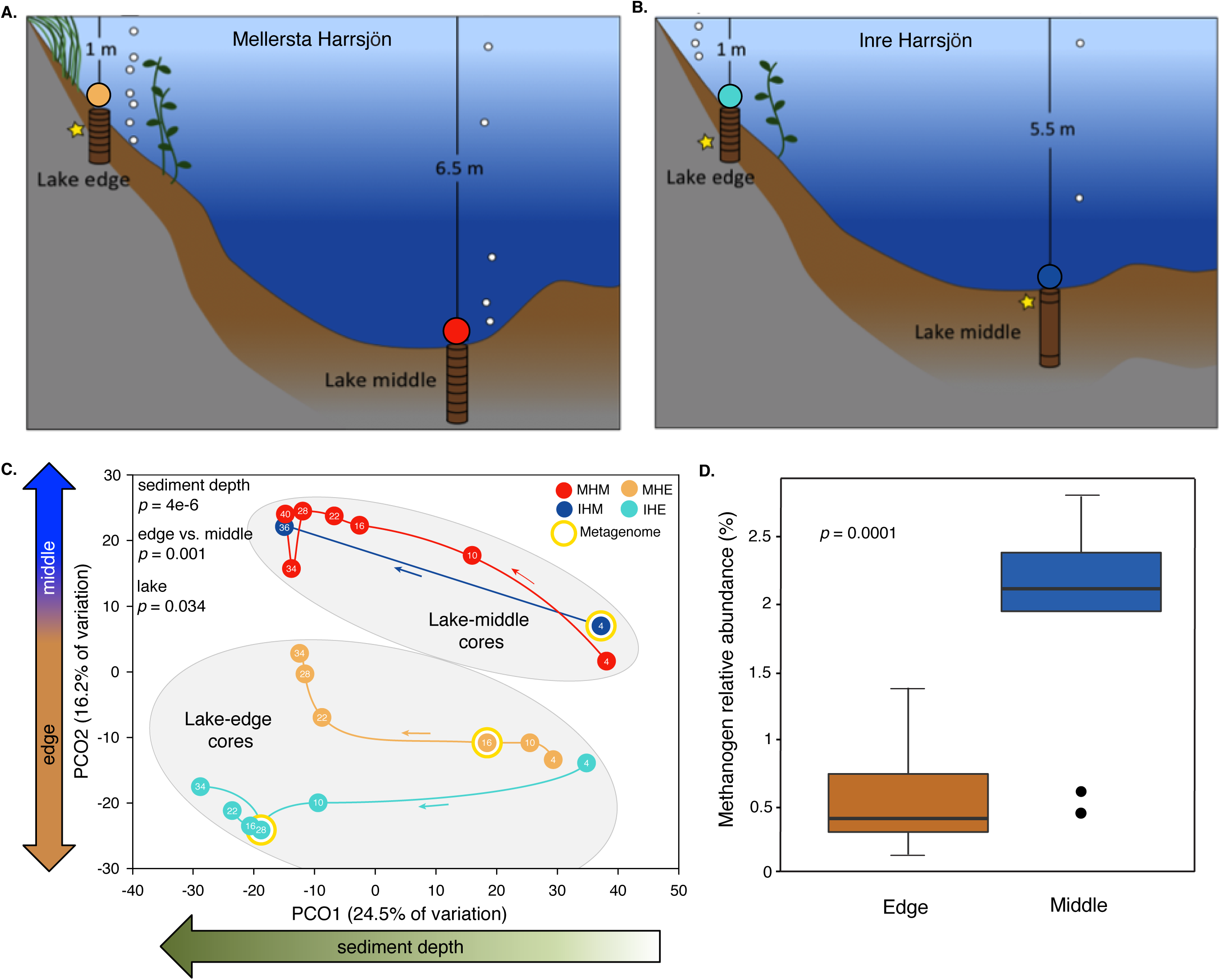
Lake sediment bacteria and archaea in two post-glacial lakes. **A, B.** Schematic overview of lakes and cores collected for DNA sequencing analyses, with core subsections indicated by horizontal lines. Cores in each lake are referred to as “Lake edge” or “Lake middle”, with overlying water depth as indicated, and the four colored circles are used to distinguish each core and/or lake location throughout the figures. Yellow stars indicate cores and depths targeted for shotgun metagenomics. **C.** Principal coordinates analysis (PCoA) of microbial community composition across samples (each core subsection, *n* = 21), based on 16S rRNA gene amplicon abundances of microbial operational taxonomic units (OTUs); circles represent samples, and samples in closer proximity have more similar microbial community composition. Thin arrows along colored lines indicate increasing depth within each core. P-values from PERMANOVA indicate how significantly microbial community composition differed according to the indicated categorical variable (significant if *p* < 0.05). **D.** Percent relative abundance of OTUs identified as methanogens in 16S rRNA gene amplicon data in lake edges compared to lake middles (P-value from Student’s T-test, significant if *p* < 0.05).

To test the relevance of these community differences to their observable CH_4_ production potential, we performed 48 *ex situ* anaerobic incubations of edge and middle sediments collected in 2012 (linked directly to our microbial and biogeochemical data) and 2013 (from the same four core locations) (Supplementary Table 9). These incubations at 5 and 22 **°**C confirmed that the lake-middle sediments had higher CH_4_ production potentials than lake-edge sediments at both temperatures (Figure 3), paralleling their higher methanogen abundances and indicating that the lake-middle methanogens can remain metabolically active at higher temperatures, despite never yet experiencing them *in situ*.

**Figure 3.**
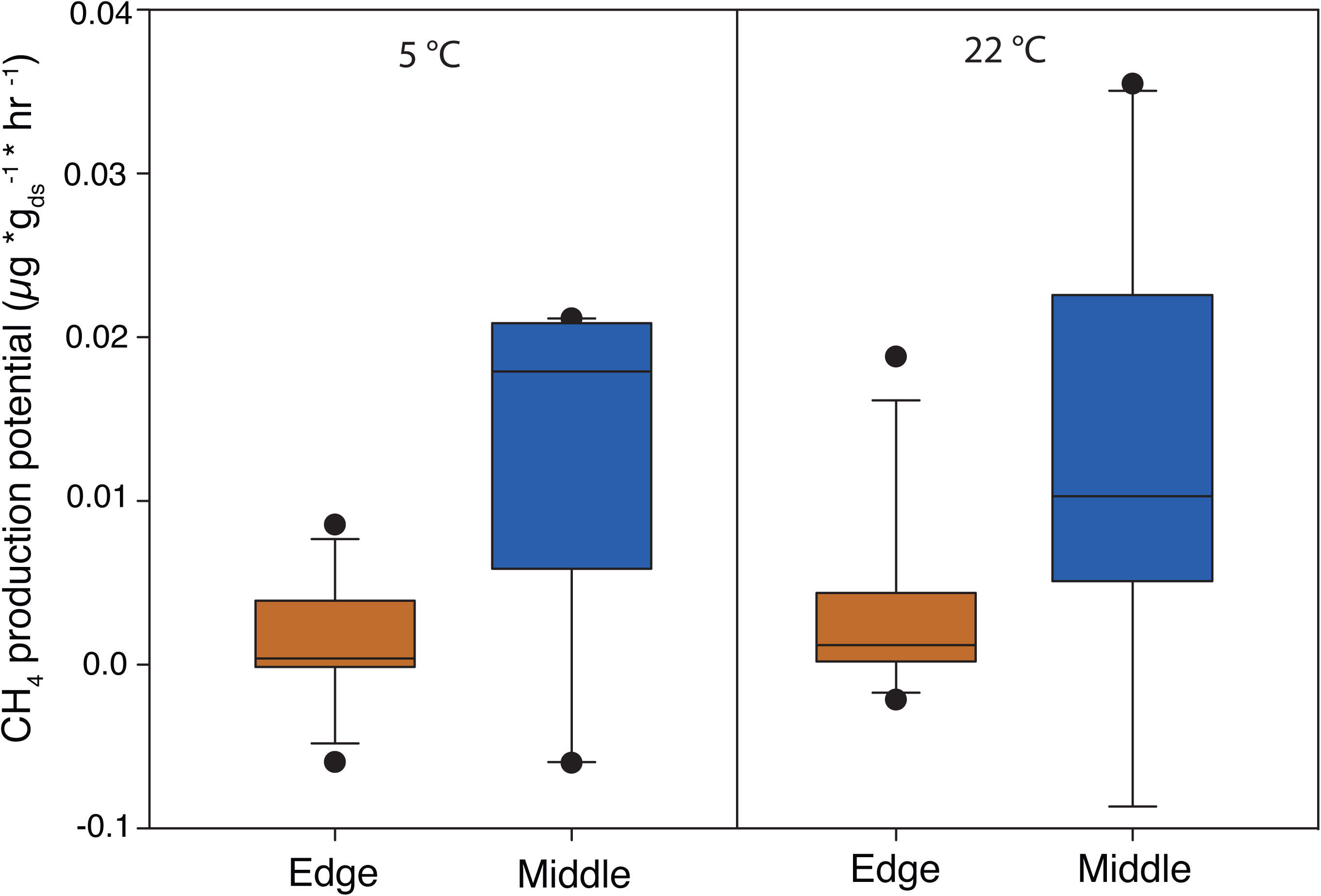
Methane production from anaerobic laboratory incubations of lake sediments. Sediments were collected from edges and middles of lakes Inre Harrsjön and Mellersta Harrsjön in 2012 and 2013 (see methods) and incubated at **A.** 5 °C (*n* = 12) and **B.** 22 °C (*n* = 12). Headspace CH_4_ concentrations were measured daily for 5 days, and average daily CH_4_ fluxes were calculated for each sample. Lines in boxes depict the median, boxes indicate 75th percentile, whiskers 95th percentile, and points are outliers. ds = dry sediment.

In order to relate microbiota from discrete depths to *in situ* CH_4_ ebullition, we partitioned ebullition to its likely source depths. We applied isotope and mass balance calculations to infer ebullitive loss (“fugitive CH_4_”) at each depth, based on stable carbon isotope values and porewater concentrations of CH_4_ and dissolved inorganic carbon (DIC) (Supplementary Table 3). From this inferred ebullitive loss, total production at each depth interval was calculated and correlated with microbiota from the same depth. Mantel tests revealed a significant correlation between microbial community composition and fugitive CH_4_ (*p* = 0.016) (Supplementary Table 5).

To more specifically investigate links between CH_4_-associated microbial functional guilds and CH_4_ chemistry, we identified multiple known CH_4_-cycling clades in the 16S rRNA gene amplicon data and applied targeted metagenomic sequencing to a subset of samples to examine diagnostic genes for CH_4_ cycling (and to assemble genomes for metabolic pathway reconstructions, discussed further below). From the metagenomes, we recovered 5,470 examples (sequencing reads) of 28 phylogenetically diverse functional genes indicative of CH_4_ production (*mcrA*) and consumption (*pmoA*) potential (Figure S5, Supplementary Table 10). We used partial least squares regressions (PLSR) and multiple linear regression (MLR) analyses to predict porewater CH_4_ concentrations from methanogen and methanotroph relative abundances, as measured via 16S rRNA gene amplicon sequencing data. When using either PLSR or MLR to predict porewater CH_4_ concentrations, a better prediction was achieved when both depth-resolved abiotic variables (*i.e*., depth, TOC, DIC, ^13^C_TOC_, S, and TOC:TS, see methods) and the relative abundances of predicted CH_4_-cycling organisms were included (PLSR: r^2^ = 0.640, *p* = 0.00001, MLR: adjusted r^2^ = 0.752, *p* = 0.0003), relative to including the abiotic variables alone (PLSR: r^2^ = 0.390, *p* = 0.002, MLR: adjusted r^2^ = 0.532, *p* = 0.0004) (Figure 4A,B, Supplementary Table 11). These results suggest that direct measurements of microbial abundances could contribute to more accurate predictions of future CH_4_ emissions, consistent with previous statistical models that have linked specific microbiota to C- and/or CH_4_-cycling dynamics in marine ecosystems and thawing permafrost peatlands^24-28^.

**Figure 4.**
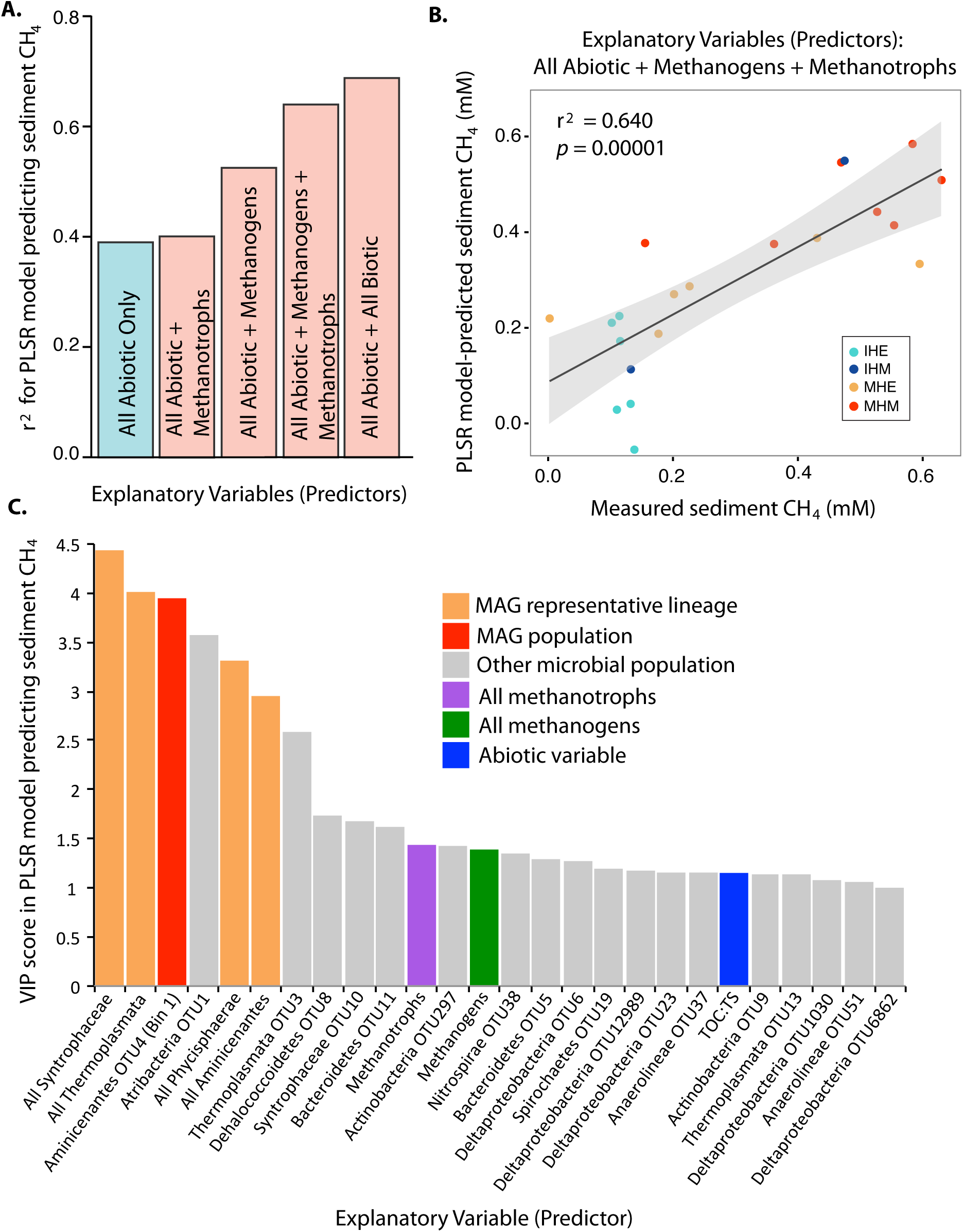
Partial Least Squares Regression (PLSR) statistical modeling to predict sediment CH_4_ concentrations. PLSR analyses tested the ability of different suites of explanatory variables to predict measured sediment CH_4_ concentrations in the four cores from 2012 across depths (*n* = 21); in all models, all measured abiotic variables (except those related to CH_4_ concentrations, see methods) were included as explanatory variables, and biotic variables were added as indicated. Biotic variables included relative abundances of specific OTUs and/or summed OTU abundances grouped by taxonomy or predicted metabolism (as indicated), from 16S rRNA gene amplicon data. **A.** Correlation coefficients (r^2^) for PLSR models predicting sediment CH_4_ using different combinations of explanatory variables. **B.** Linear regression of measured and model-predicted sediment CH4, considering all abiotic variables and methanogen and methanotroph abundances as explanatory variables; each point is a sample, colored by core. **C.** For the model with the highest r^2^ (rightmost in panel A), VIP scores are plotted to indicate the relative contribution of each explanatory variable; a VIP score > 1 is considered significant, and higher VIP scores indicate a more significant contribution to the model; all VIP scores > 1 are shown (*n* = 26 out of *n* = 153 total, Supplementary Table 14).

By expanding our PLSR analyses to consider the full microbial community, in addition to known CH_4_-cyclers, our ability to predict CH_4_ concentrations improved further. This analysis considered the following groupings of 16S rRNA gene abundances as explanatory variables for the prediction of porewater CH_4_ concentrations: 1) each operational taxonomic unit (OTU) at > 1 % relative abundance in any sample (Supplementary Table 4), 2) summed lineage abundances of all bacteria and archaea (mostly at the phylum or class levels, see Figure S3 for groupings), and 3) summed abundances of the most highly resolved lineage representative in the amplicon data for each metagenome-assembled genome (MAG, a population genome computationally reconstructed from shotgun metagenomic community DNA sequencing data, Supplementary Table 12). In two cases, a MAG was linked directly to a specific OTU in the amplicon data through a co-binned 16S rRNA gene sequence in the MAG, such that the MAG relative abundance could be inferred from the amplicon data. In all other cases, the summed abundances of amplicon OTUs in the same lineage as the MAG were used as proxies for MAG abundances.

Four of the top five microbial groups most predictive of porewater CH_4_ concentrations in the PLSR analyses were lineages for which we were able to reconstruct a MAG (Figure 4C, Supplementary Tables 13-14), thus organization into MAGs helped to unravel the specific metabolic processes most predictive of carbon chemistry. In total, five MAGs were reconstructed with > 85 % completeness and < 6 % contamination (Supplementary Discussion). The best overall predictor of porewater CH_4_ concentrations was the Syntrophaceae class of Deltaproteobacteria, which was considered in the PLSR analysis as the summed abundance of all OTUs in this clade. Syntrophaceae are known to be syntrophic (obligately mutualistic) with methanogens and produce the hydrogen needed for methanogenesis^29^. Consistent with hydrogen production, the Syntrophaceae MAG revealed 15 hydrogenase-associated genes, along with the capacity to ferment diverse carbon compounds (particularly carbon-sulfur compounds), with the added potential capacity for respiration (see Supplementary Discussion). Though the Syntrophaceae were overall most predictive of porewater CH_4_ concentrations, the most significantly predicitive single OTU was a member of the candidate phylum Aminicenantes, which we also recovered as a MAG. While this lineage has been previously predicted to be fermentative, saccharolytic, and/or aerobic^30-32^, our lake sediment genome revealed metabolic potential for several C1 metabolic processes, including methylotrophy through the assimilation of methylamines, methane-thiols, and/or dimethylsulfide, similar to previous recoveries of complete Wood-Ljungdahl pathways for C1 metabolism via carbonyl and methyl pathways in this lineage^33^. The predicted capacity for methylotrophy could explain the strong correlation between Aminicenantes relative abundance and porewater CH_4_ concentrations.

The relative abundances of two other lineages with MAGs, the Thermoplasmata (a group of Archaea) and Phycisphaerae (a class of Planctomycetes bacteria), were also strongly predictive of both porewater CH_4_ concentrations in the PLSR analysis and of calculated fugitive CH_4_ in linear regressions (Supplementary Tables 14-15). Phylogenetic analyses showed that the Thermoplasmata MAG was derived from a divergent member of the Thermoplasmatales order, and it encodes the capacity for CO_2_ production from formate, along with peptide and amino acid degradation (as previously indicated^34^) and complex carbon degradation. Our recovered Phycisphaerae population genome appears to have the capacity to metabolize a wide variety of complex carbon compounds, potentially via fermentation, consistent with previous predictions for the Planctomycetes phylum^35^. While direct ties to CH_4_ are not obvious in these two genomes, we speculate that their contributions to overall carbon cycling may be driving these strong correlations with CH_4_ concentrations and emissions.

Interestingly, the only lineage represented by a MAG that was not a significant predictor of porewater CH_4_ concentrations in the PLSR analysis was a member of the archaeal Methanomassiliicoccales, a lineage previously presumed to consist exclusively of obligate H_2_-dependent methylotrophic methanogens^36,37^. While we cannot make a definitive claim based on a single MAG, we hypothesize that our lake sediment Methanomassiliicoccales population does not have the capacity for methanogenesis, as we did not recover any genes from the methanogenesis pathway in this 95% complete genome. The genome does encode a complete pathway for propionate fermentation and partial pathways that may be indicative of the potential to ferment benzoate, butyrate, and succinate.

In conclusion, we found significant differences in the slope of the temperature vs. CH_4_ flux relationship between sub-arctic lake edges and middles, suggesting that radiative forcing (temperature) and a concomitant increase in microbial metabolic rates are not the only primary controls on CH_4_ emissions. Significant differences in microbial community composition between lake edges and middles, including significantly higher methanogen abundances in lake middles, and significantly higher CH_4_ emissions from lake middle sediments when incubated at the same temperatures as lake edges suggest that sediment microbial community composition contributes to spatial differences in the response of CH_4_ emissions to increasing temperature. In addition, the abundances of CH_4_-cycling organisms and their reconstructed population genomes (MAGs) were significantly better predictors of sediment CH_4_ concentrations than abiotic variables alone. Syntrophic lineages, which can generate the hydrogen required for hydrogenotrophic methanogenesis, and lineages capable of C degradation to CO_2_ (also potentially ‘upstream’ of methanogenesis) were also predictive of sediment CH_4_ concentrations. Together, these results suggest that when lake middles reach the temperatures of lake edges, they may emit even more CH_4_ than the lake edges currently do, such that our projected future CH_4_ emissions may be underestimating contributions from subarctic lakes, and that knowledge of microbial community composition and metabolism could improve these predictions. Future investigations that consider the combined effects of microbiota, carbon quality, and temperature on lake CH_4_ emissions will help to provide a more comprehensive understanding of spatiotemporal controls on global CH_4_ emissions.

## Methods

### Field site and sample collection

Stordalen Mire is a subarctic peatland complex located 10 km east of Abisko in northern Sweden (68°21′N, 19°02′E). Lakes Mellersta Harrsjön and Inre Harrsjön are 1.1 and 2.3 ha in area, reaching maximum depths of 7 and 5 m, respectively^38^. These lakes are post-glacially formed. Mellersta Harrsjön receives water from a small stream while Inre Harrsjön is fed through groundwater and runoff from the surrounding mire. Ebullitive and diffusion-limited CH_4_ emissions from these lakes have been documented, using floating funnels and chambers distributed across the lakes and sampled frequently^2,9,12^. Ebullition varies spatially with higher emissions from shallow zones and in the presence of plants^9,15^.

We collected quadruplicate sediment cores (four cores from two locations in each of two lakes: Mellersta Harrsjön edge (68°357832’N, 19°042046’E) and middle (68°358291’N, 19°042132’E) and Inre Harrsjön edge (68°357880’N, 19°048525’E) and middle (68°358418’N, 19°045650’E)) on July 10 and 18, 2012 at the Stordalen Mire nature reserve, a research site near Abisko, northern Sweden (Supplementary Table 1). Samples were taken from cores (as described below) along a depth gradient (ranging from 4 - 40 cm) for geochemical measurements and microbial DNA sequencing data.

### Geochemical data collection and analysis

For each set of four cores, we sampled the first core for sediment C, N, and S (weight percent), percent total organic carbon, and bulk sediment ^13^C_TOC_ and ^15^N_TOC_. Samples of 1 cm^3^ were taken in 6 cm increments from the top of the core to the bottom. The samples were then dried, ground, and split into an untreated sample for total carbon (C) and an acidified TOC sample. Details regarding sample preparation for measurement on a Perkin Elmer 2400 Series II CHNS/O Elemental Analyzer at the University of New Hampshire (UNH) were described previously^15^. Repeatability error was established by analyzing replicate samples and calculating the standard deviation. Duplicate samples were run approximately every 10 samples. Potential outliers were also run in duplicate. Isotopic analysis was performed by combusting dried sediment samples in a Costech ECS 4010 elemental analyzer coupled to a Thermo Trace GC Ultra isotope ratio mass spectrometer (IRMS), based on calibration with acetanilide, Atlantic cod, black spruce needles, sorghum flour, corn gluten, NIST 1515 apple leaves and tuna muscle standards (UNH Stable Isotope Lab). In 2013 we also collected sediment cores in the same locations in these lakes. We report sediment textural analyses from these cores as % sand, % silt, and % clay (Supplementary Table 3). Those samples were dried and run through a laser particle size analyzer (Malvern Mastersizer 2000).

The second replicate core was used for quantifying total CH_4_ in the core sediment reported in μM. After coring, we pulled 2 cm^3^ sediment plugs using cut plastic syringes through pre-drilled holes cut at 4 cm increments along the core liner. The sediment plugs were transferred to 30 ml serum vials containing 5 ml of 2 M NaOH, capped quickly and shaken^39,40^. After sitting overnight then heating for 1 hour at 60 **°**C, the headspace of the vials was analyzed for CH_4_ using a Shimadzu GC-2014 gas chromatograph with a flame ionizing detector^9^. The CH_4_ measured represents the total, that is, nearly all of the CH_4_ dissolved in the water from the sediment plug and any bubbles that may have been trapped in the sediment. The remaining sediment samples in the vials were weighed and dried to constant weight to determine the mass of water in the samples to be used for calculating the CH_4_ concentration in μM.

The third replicate core was used for measurement of DIC. Rhizon samplers were inserted every 2 cm through pre-drilled holes in the core and a vacuum was pulled with a 30 ml polypropylene syringe. The first ∼1 ml of sediment water was discarded because of contamination with DI water. After 10 ml of sediment pore water was collected, it was injected to a 30 ml evacuated serum vial with 1 ml 30% H_4_PO_4_ solution. This caused forms of inorganic C in the water to form CO_2_. A headspace sample was then extracted and run on an infrared gas analyzer (IRGA) to determine the CO_2_ concentration.

Methods for measuring ebullition and water temperature have been described previously^9^. In brief, measurements of CH_4_ bubble flux during the ice-free season (June to September) have been ongoing at these lakes since 2009. A total of 40 bubble traps, distributed in a depth-stratified sampling scheme were sampled frequently (every 1-3 days). For this study, averages of CH_4_ bubble flux were calculated for each lake by binning data from edge and middle areas separately in 1°C intervals (total of 4-22°C) of corresponding surface sediment temperature. For this we used flux and temperature data collected from 2009-2014. Water and surface sediment temperatures were measured in profiles continuously using intercalibrated Onset HOBO v22 loggers, as previously described^9^ (data are available here: https://bolin.su.se/data/). The binned flux data were used to construct Arrhenius equations in order to investigate differences in temperature response on the ebullition from edge and middle areas.

Porewater isotopic composition was determined in samples from cores collected in the same locations in 2014. Methods were described previously^24^. Briefly, sample vials that were collected for CH_4_ and dissolved inorganic carbon (DIC) were acidified with 0.5 ml of 21% H_3_PO_4_ and brought to atmospheric pressure with helium. The sample headspace was analyzed for d13C of CH_4_ and CO_2_ on a continuous-flow Hewlett-Packard 5890 gas chromatograph (Agilent Technologies) at 40°C coupled to a FinniganMAT Delta S isotope ratio mass spectrometer via a Conflo IV interface system (Thermo Scientific).

### DNA extraction and 16S rRNA gene sequencing

A fourth replicate core was collected for DNA extraction. After coring, we pulled 2 cm^3^ sediment plugs using cut plastic syringes through pre-drilled holes cut at 4 cm increments along the core liner. Samples were immediately put in Eppendorf tubes and placed in a cooler until returned to the research station where they were stored at −80 °C until extraction.

For DNA extraction from each core depth range, 0.25 g of sediment was collected under sterile conditions and added to a MoBio PowerSoil DNA Isolation Kit (MoBio, Inc., Carlsbad, CA, USA). DNA was extracted according to the manufacturer’s instructions. PCR amplification and sequencing were performed at the Environmental Sample Preparation and Sequencing Facility (ESPSF) at Argonne National Laboratory, in accordance with previously described protocols^41-43^. Briefly, 515F and barcoded 806R primers with Illumina flowcell adapter sequences were used to amplify the V4 region of bacterial and archaeal 16S rRNA genes^44^. Each 25 µl PCR reaction contained 12 µl of PCR water (MoBio, Inc., Carlsbad, CA, USA), 10 µl of 1x 5 PRIME Hot Master Mix (5 PRIME Inc., Bethesda, MD, USA), 1 µl each of F and R primers (5 µM concentration, 200 pM final), and 1 µl of template DNA. PCR cycling conditions were as follows: 94 °C for 3 min, 35 cycles of [94 °C for 45 s, 50 °C for 60 s, and 72 °C for 90 s], 72 °C for 10 min. A PicoGreen assay (Life Technologies, Grand Island, NY, USA) was used to measure amplicon concentrations. Equimolar concentrations for each barcoded sample were combined and then cleaned with the UltraClean PCR Clean-Up Kit (MoBio Inc., Carlsbad, CA, USA) and then quantified using the Qubit (Invitrogen, Carlsbad, CA, USA). The pool was then diluted to 2 nM, denatured, and then diluted to a final concentration of 4 pM with a 10% PhiX spike for sequencing on the Illumina MiSeq platform.

### Quantitative PCR (qPCR)

A quantitative polymerase chain reaction (qPCR) was performed to measure microbial abundances in units of 16S rRNA gene copies per g wet sediment^43,45^. Each reaction used 5 µl of 2X SYBR Green PCR Master Mix (Applied Biosystems, Carlsbad, CA, USA), 4 µl of template DNA, and 1 µl of primer mix. The 16S rRNA gene 1406F/1525R primer set (0.4 µM, F - GYACWCACCGCCCGT and R - AAGGAGGTGWTCCARCC) was designed to amplify bacterial and archaeal 16S rRNA genes. The rpsL primer pair (0.2 µM, F - GTAAAGTATGCCGTGTTCGT and R - AGCCTGCTTACGGTCTTTA) was used for inhibition control samples to amplify *Escherichia coli* DH10B only. Three dilutions (1/100, 1/500, and 1/1000), as well as an inhibition control (1/100 dilution of *E. coli* DH10B genomic DNA spiked into a 1/100 dilution of the sample), were run in triplicate for each sample and standard. For the standards, *E. coli* DH10B genomic DNA dilutions of 10^−2^, 10^−3^, 10^−4^, 10^−5^ and 10^−6^ of the 20 ng/µl stock solution were used. The qPCRs were run on the ViiA7 Real-Time PCR System (Applied Biosystems, Carlsbad, CA, USA), with cycling conditions as follows: 10 min at 95 **°**C, 40 cycles of [15 s at 95 **°**C, then 20 s at 55 **°**C, then 30 s at 72 **°**C]. A melt curve was produced by running a cycle of 2 min at 95 **°**C and a final cycle of 15 s at 60 **°**C. The cycle threshold (Ct) values were recorded and analyzed using ViiA7 v1.2 software, and 16S rRNA gene copy numbers were calculated for each sample, accounting for the genome size (4,686,137 bp) and 16S rRNA gene copy number (7) of the standard.

### Incubations for CH_4_ production rates

Anaerobic incubations of lake sediment samples were performed to assess rates of production of CH_4_. Four replicate sediment samples (4 ml) from three depths in 2012 (0-5, 10, 20 cm) were collected in the field and immediately sealed in a 120 ml serum vial. The headspace was flushed for 5 minutes with UHP N_2_ to establish an anaerobic headspace. The vials were stored in coolers, taken to the research station, and then stored as follows: 2 vials were incubated at 5°C and 2 vials were held at room temperature (22°C) for each depth. Five ml of headspace was sampled daily for five days and analyzed on a Flame Ionization Gas Chromatograph (GC) to determine CH_4_ fluxes. Fluxes were normalized by sediment mass after incubations when vials were dried and weighed to determine sediment dry weight. We also report data from incubations in 2013 that were run the same way with samples collected at depths consistent with changes in core sediment transitions: Inre Harrsjön edge: 2.5, 27.5, 47.5 cm; Inre Harrsjön middle: 4.5, 35, 60 cm; Mellersta Harrsjön edge: 7.5, 22.5, 37.5 cm; and Mellersta Harrsjön middle: 2.5, 27.5, 47.5 cm.

### Calculations of depth-resolved fugitive CH_4_

Depth-resolved fugitive CH_4_ (CH_4_ released from the sediments) was calculated from concentration and stable carbon isotopic composition of CH_4_ and DIC in sediment porewater^46^. The approach leverages that fact that 1) microbial fermentation and respiration, which generate CO_2_, do not fractionate carbon, while methanogenesis, which generates CH_4_ and CO_2_ (1:1), does fractionate carbon, and 2) that DIC largely remains dissolved in water while dissolved CH_4_ escapes porewater by ebullition. In this framework, the measured isotopic composition of CH_4_ in porewater was used to calculate the fraction factor associated with methanogenesis, assuming the starting isotopic composition of the substrate matched that measured for organic carbon in the sediment. This fractionation factor, along with the measured isotopic composition of DIC in porewater, was used to determine the relative amount of DIC that came from methanogeneis versus non-fractionating pathways (*e.g*., fermentation). Because any CO_2_ produced was assumed to stay dissolved in porewater, the relative amount of DIC generated from methanogenesis could be multiplied by the measured concentration of DIC to determine the concentration of CO_2_ and CH_4_ generated through methanogenesis. This generated CH_4_ concentration was larger than the actual measured concentration of CH_4_ in porewater, and the difference between the two was assigned as ‘fugitive’ methane. Calculations assumed that the system was at steady state.

### 16S rRNA gene sequence processing and OTU table generation for microbial analyses

Sequences were processed as previously described^43^. Briefly, after demultiplexing by sample, each pair of forward and reverse 16S rRNA gene reads was merged. Sequences were then quality-filtered, and singletons were removed with QIIME^47^ and UPARSE^48^. Dereplicated sequences were then clustered at 97% nucleotide identity using UCLUST v7^49^ to generate a database containing one sequence for each operational taxonomic unit (OTU). Sequencing reads from the full dataset were then clustered to the database to generate an OTU table. Each OTU was assigned taxonomy via the Ribosomal Database Project taxonomic classifier^50^, and all OTUs assigned as mitochondria or chloroplasts were removed. The resulting OTU table was rarefied to 3,000 16S rRNA gene sequences per sample. Following this OTU table curation, 36 samples across 21 core-depth combinations were retained, of which 30 were replicates (*i.e*., 15 pairs). For each pair of replicates, each OTU count was averaged (for 14 of 15 pairs, replicates were indistinguishable, Figure S6), and the averages were used for all downstream analyses. For the six samples without successful replicates, OTU counts from a single sample were used.

### Metagenomic sequencing, genome reconstruction and annotation, and methane-cycling functional gene characterization

Based on preliminary 16S rRNA gene amplicon sequencing data from 8 samples (IHM4, IHM36, IHE4, IHE28, MHM4, MHM34, MHE4, and MHE16), three samples with the most distinct microbial communities (IHM4, IHE28, and MHE16) were selected for metagenomic sequencing to maximize recovery of diverse microbial populations. DNA (from the same extractions described above for 16S rRNA gene sequencing) was sent to the Australian Centre for Ecogenomics for metagenomic library construction and sequencing on the Illumina NextSeq platform, as previously described^25,26^. Metagenomic assembly, genome binning to recover microbial metagenome-assembled genomes (MAGs), and annotation (to predict gene functions and reconstruct metabolic pathways) were performed as previously described^51^. Briefly, each metagenome was separately assembled using the CLC *de novo* assembler v4.4.1 (CLCBio, Denmark), reads were mapped to contigs using BWA v0.7.12-r1039^52^, and the mean coverage of contigs was obtained using the ‘coverage’ command of CheckM v1.0.6^53^. Genomes were binned using MetaBAT v0.26.3^54^ with all five preset parameters (verysensitive, sensitive, specific, veryspecific, superspecific), and genome completeness and contamination were estimated using CheckM^53^. To investigate predicted metabolic functions of interest in the metagenomic data, metagenomic reads with sequence similarity to genes diagnostic of specific metabolic functions (*e.g*., methane monooxygenase, *pmoA*, and methyl-coenzyme M reductase, *mcrA*, indicative of aerobic methane oxidation and methanogenesis, respectively) were identified using GraftM^55^.

### Sequencing data availability

Data are currently available here: https://isogenie-db.asc.ohio-state.edu/datasources#lake_data. Upon publication, sequencing data from this study will be available at NCBI, with accession numbers provided here.

### Statistical analyses

Unless otherwise indicated, statistical analyses were performed using PRIMER v7^56,57^. The rarefied OTU table was square-root transformed, and Bray-Curtis similarity matrices were generated for sample comparisons and used to make a Principal Coordinates Analysis (PCoA) plot. We used permutational ANOVA (PERMANOVA) to test for significant differences in microbial community composition between categorical groups of samples (*e.g*., between the two lakes and between the edges and middles of the lakes), and we used Mantel tests with Spearman’s rank correlations to compare microbial community composition (Bray-Curtis similarity matrices) to continuous variables (Euclidean distance matrices), including sediment depth and biogeochemical data. ANOVA and linear regression analyses (Supplementary Tables 8 and 15) were performed with StatPlus v6.1.7.0.

We performed partial least squares regressions (PLSR) in the R programming language via the package PLS (function PLSR)^58-60^ to predict measured sediment CH_4_ concentrations from biotic and abiotic variables, similar to our previously described PLSR analyses^25^. Briefly, PLSR models a causal relationship between explanatory variable(s) (in this case, abundances of abiotic measurements and/or microorganisms) and the response variable being predicted (here, measured sediment CH_4_ concentrations). Abiotic variables included all depth-resolved abiotic measurements that were not directly related to CH_4_, as such measurements could be confounding variables in our analysis. The included abiotic variables were: depth, TOC, ^13^C_TOC_, DIC, S, and TOC:TS. The PLSR analysis yielded Pearson’s product moment correlations between measured environmental and/or geochemical variables, the abundances of microbial lineages, and the abundances of specific microbial populations, allowing for a quantification of the added value of microbial abundances in predicting sediment CH_4_ concentrations, relative to predictions from abiotic factors alone. Variance in projection (VIP) scores for each explanatory variable indicate the extent to which that variable was predictive of the response variable (*i.e*., sediment CH_4_ concentrations), with VIP scores ≥ 1 considered to be highly significant^61^.

## Acknowledgements

We would like to acknowledge the following funding in support of this project: the Northern Ecosystems Research for Undergraduates program (NERU; National Science Foundation REU site EAR-1063037, PI Varner), a U.S. National Science Foundation MacroSystems Biology grant (NSF EF #1241037, PI Varner), U.S. Department of Energy grants (DE-SC0010580 and DE-SC0016440, Co-lead PI Rich; DE-SC0010338 and DE-SC0019063, PI Neumann), the Swedish Research Council (VR) with grants to P. Crill (2007-4547 and 2013-5562). Thanks to staff at the Polar Research Secretariat’s Abisko Research Station (ANS). Thanks to Kaitlyn Steele, Florencia Fahnestock, Kiley Remiszewski, Carmody McCalley, and NERU participants Sophia Burke, Joel DeStasio, Lance Erickson, and Madison Halloran for assistance in sample collection and analysis, and Jacob Setera and Steve Phillips (UNH) for assistance with the CHNS elemental analysis.

