## Supplementary Text and Figures for "Diverse Arctic lake sediment microbiota shape methane emission temperature sensitivity"

### Supplementary Discussion

*Predicted metabolisms and abundance profiles of population genome bins*

The Bin 1 metagenome-assembled genome (MAG), from Candidate Phylum Aminicenantes (previously predicted to be fermentative, saccharolytic, and/or aerobic[1]), is predicted to have the capacity for several C1 metabolic processes, including methylotrophy through the assimilation of methylamines, methane-thiols, and/or DMS, and possibly $CO_2$ fixation through a near-complete Wood-Ljungdahl pathway (but the direction of the WLP is unclear – it may operate in the reverse direction). The dominant Aminicenantes population is also predicted to have a complete pathway for propionate fermentation. The 16S rRNA gene data suggest that the Aminicenantes lineage generally increases in abundance with depth in all four cores, and the population from which the genome was recovered (the most abundant Aminicenantes OTU) is particularly abundant in the deeper regions of the Inre Edge core. Across the three metagenomes, Bin 1 has very high coverage in the Inre Edge 28 cm sample, near 0x coverage in the Inre Middle 4 cm core and very low coverage in the Mellersta Edge 16 cm sample.

The Bin 16 MAG is taxonomically classified as a member of the archaeal Methanomassiliicoccaceae, presumed to be a methanogenic lineage, with some populations capable of fermentation in addition to methanogenesis[2]. At 95.2% complete, 6 of the 7 enzymes in the methanogenesis pathway, including all subunits of the methyl coenzyme M reductase (MCR) complex for generating methane, were not detected in this MAG, suggesting that this population may lack the capacity for methanogenesis. However, with a single representative genome for this lineage and a complete lack of mcrA genes recovered in the entire assembled dataset (including all bins and unbinned contigs, despite clear evidence that methanogenesis occurs in this system), we are reluctant to do more than suggest that there is some possibility that this lineage might not be methanogenic. Instead, the dominant Methanomassiliicoccaceae genome has a complete pathway for propionate fermentation and partial pathways suggestive of the potential to ferment other compounds, potentially including benzoate, butyrate, and succinate. It also encodes clostripain for extracellular peptide degradation. The 16S rRNA gene data suggest that the Methanomasillicoccaceae (as a lineage) generally maintain similar abundances or increase in abundance with depth in the four cores, and the specific population from which the MAG was recovered (the most abundant

Methanomasilliicoccaceae OTU) generally increases in abundance with depth and is particularly abundant in the Inre Edge core. Across the three metagenomes, Bin 16 is at moderate (average 18x) coverage depth in the Inre Edge 28 cm sample and at near 0x coverage in both the Inre Middle 4 cm and Mellersta Edge 16 cm samples.

The Bin 19 MAG is classified as a divergent member of the archaeal Thermoplasmatales[3]. Of the predicted proteins encoded by the genome, 38% are hypothetical, meaning that no known homologs with classified functions were recovered in our analyses, suggesting a relatively high amount of previously unknown or divergent functional capacity. The genome indicates the capacity for $CO_2$ production from formate and amino acid degradation (*e.g.*, complete pathways for L-arginine and L-citrulline degradation to ammonium and $CO_2$, along with L-histidine degradation). It also encodes gingipain for extracellular peptide degradation, along with genes involved in complex carbon degradation, including beta-xylosidase, alpha-amylases, and chitodextrinase, and it has the capacity for glycogen storage and breakdown. Although the MAG could not be directly linked to a 16S rRNA gene sequence, the 16S rRNA gene data for the Thermoplasmatales lineage suggest that these organisms generally increase in abundance with depth. Across the three metagenomes, Bin 19 is at approximately 0x coverage in the Inre Edge 28 cm and Inre Deep 4 cm samples and at moderate abundance (average 17x coverage) in the Mellersta Edge 16 cm sample.

Consistent with previous predictions for the Planctomycetes lineage[4], the Phycisphaerae MAG (Bin 34) appears to have the capacity to metabolize a wide variety of complex carbon compounds. Genes encoding the following enzymes were recovered, supporting diverse carbon degradation capacities: trehalose utilization, beta-xylosidases, a variety of glycosyl hydrolases, pectate lyases, beta-xylanase, chitinase, beta-lactamases, cellulose synthase, beta-hexosaminidase, xanthine dehydrogenase, gellan lyase, xylonate dehydratase, beta-galactosidases, alpha-L-fucosidase, alpha-L-arabinofuranosidases, cellobiose-2-epimerases, cellulases, and exo-beta-D-glucosaminidase. With 41% hypothetical proteins, specific metabolic processes are difficult to infer, but the genome also encodes multiple hydrogenases and evidence for C1 metabolism of methylamines, possibly from choline degradation, in addition to some evidence for nitrogen fixation (as previously reported[5]) and sulfur metabolism. Across the three metagenomes, Bin 34 is at moderate abundance (average 12x coverage) in the Inre Edge 28 cm sample and at near 0x coverage in the Inre Middle 4 cm and Mellersta Edge 16 cm samples.

Bin 41, a member of the Syntrophaceae lineage of Deltaproteobacteria, is hypothesized to be a fermenter of diverse carbon compounds, particularly carbon-sulfur compounds, and may have the capacity for respiration (*e.g.*, a predicted fermentation/respiration switch protein was encoded in the genome). Many amino acid transporters and a near-complete pathway for pyruvate fermentation were encoded. Bin 41 was at moderate (average 12x) coverage in the Inre Middle 4 cm sample and near 0x coverage in both the Inre Edge 28 cm and Mellersta Edge 16 cm samples.

Figure S1

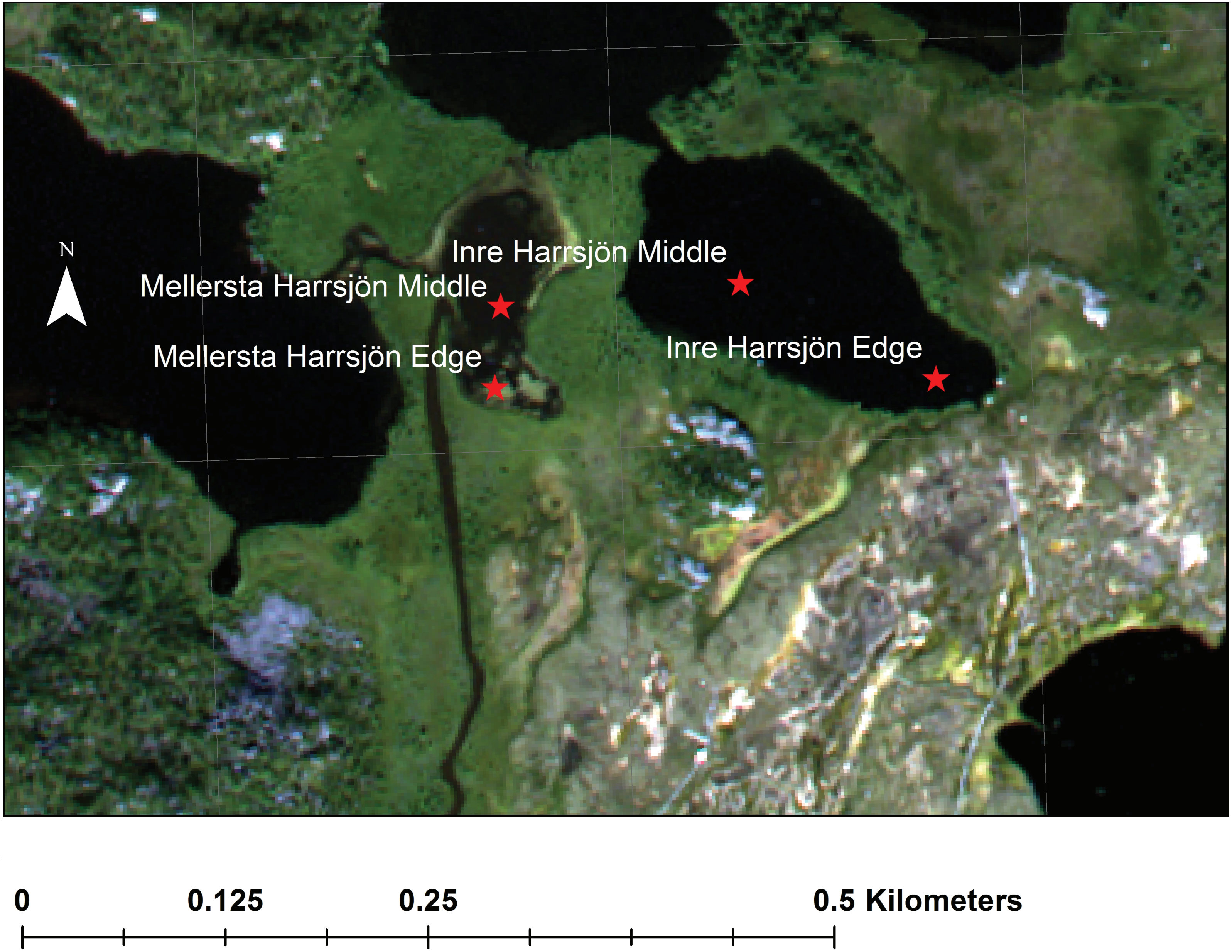


**Supplementary Figure 1. Satellite image of 2012 core sampling locations.**

Figure S2

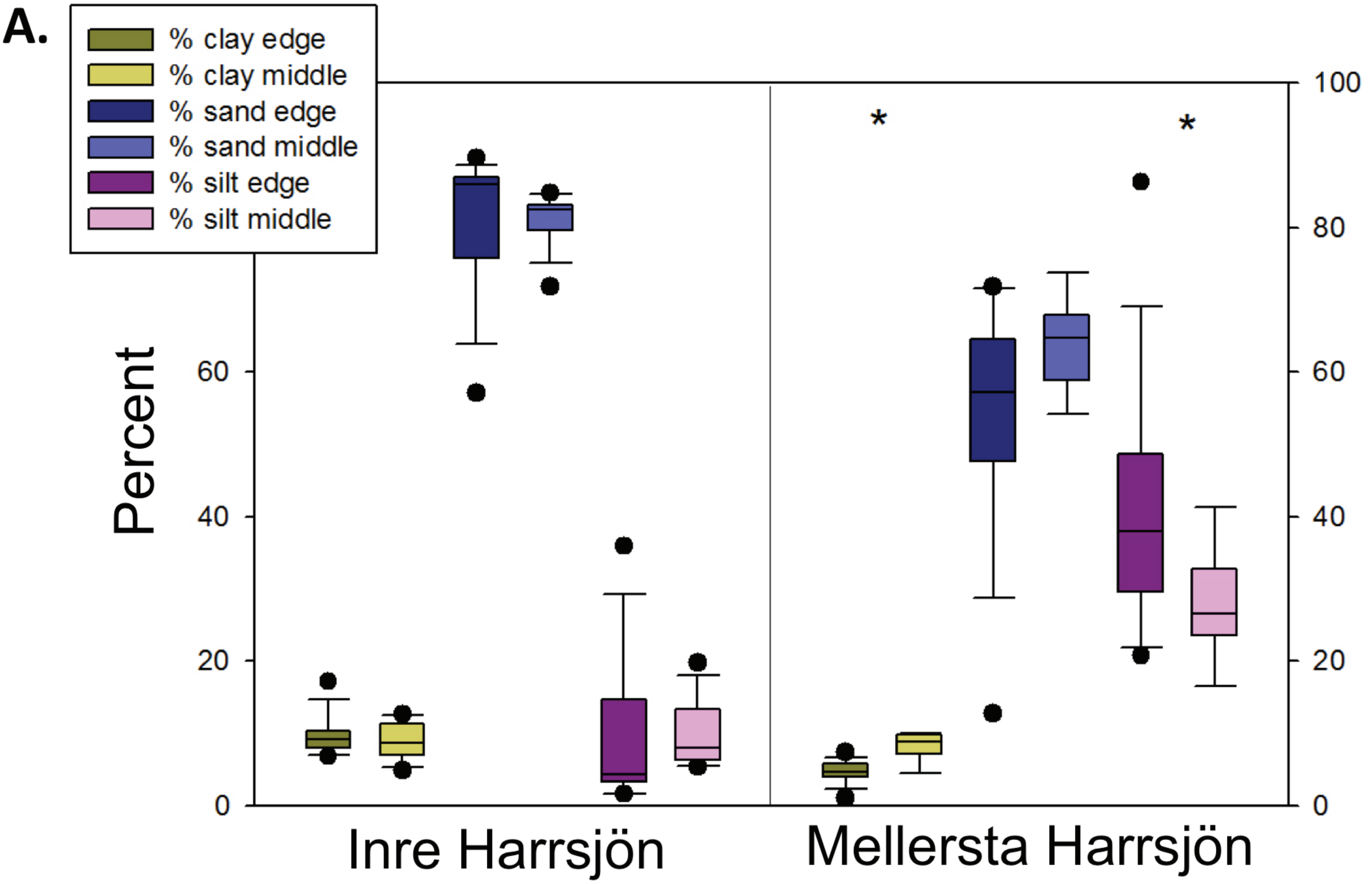


**B.**

| Lake edge vs. middle | clay | sand | silt |
|---|---|---|---|
| Inre Harrsjon | 0.412 | 0.9978 | 0.7912 |
| Mellersta Harrsjon | **<0.0001** | 0.1031 | **0.0425** |

**Supplementary Figure 2. Texture analysis in edges vs. middles of each lake.** **A.** Sand, silt, and clay percentages for each lake, edge vs. middle. Asterisks indicate significant edge vs. middle differences. **B.** Tukey Kramer T-test of differences in sand, silt, and clay in edges vs. middles. Bold indicates significant differences.

Figure S3

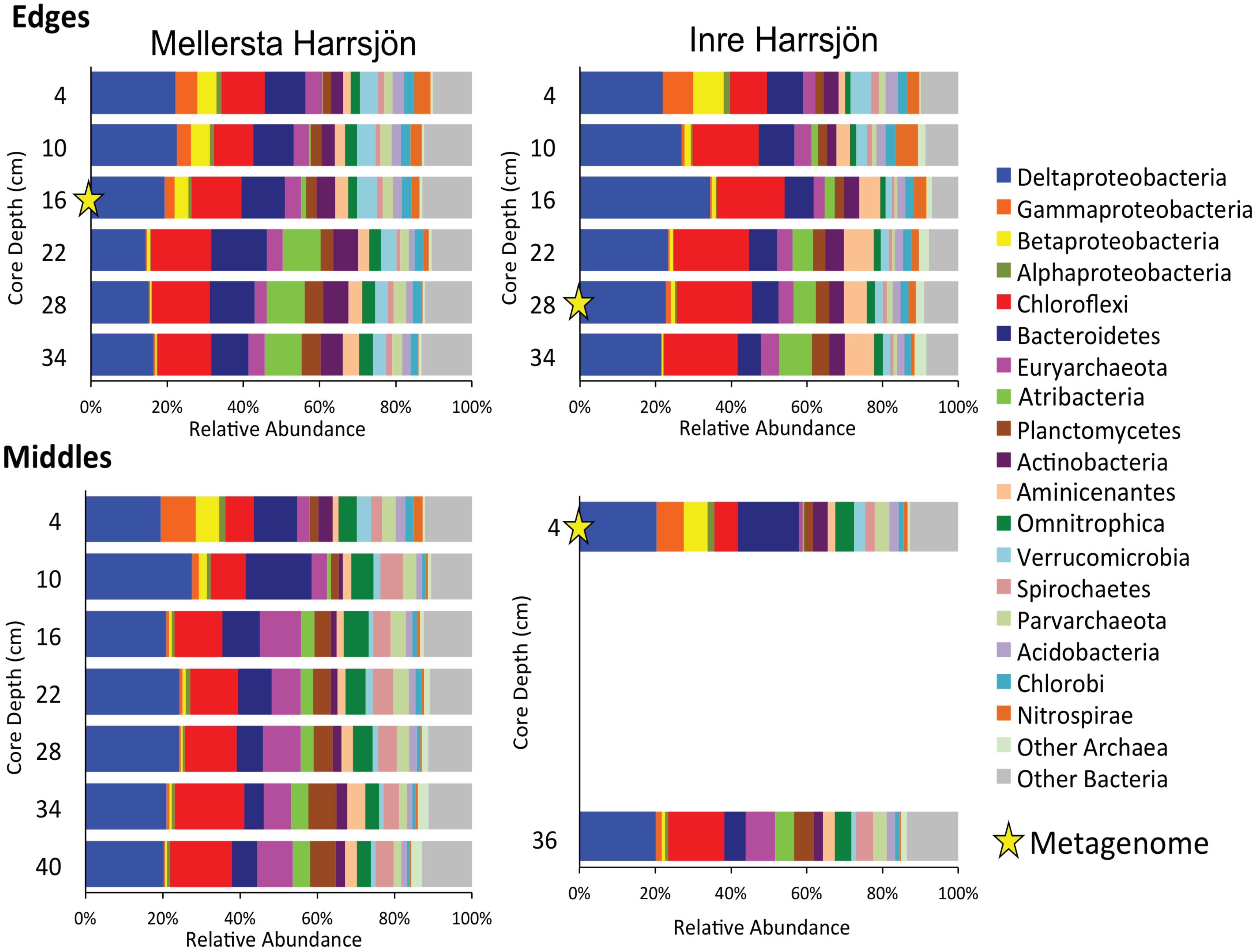


**Supplementary Figure 3. Relative abundances of microbial taxa by core and sediment depth**. Microbial relative abundances in 16S rRNA gene amplicon sequencing data (Bacteria grouped by phylum, or class for Proteobacteria; Archaea grouped together, see Supplementary Table 5).

Figure S4

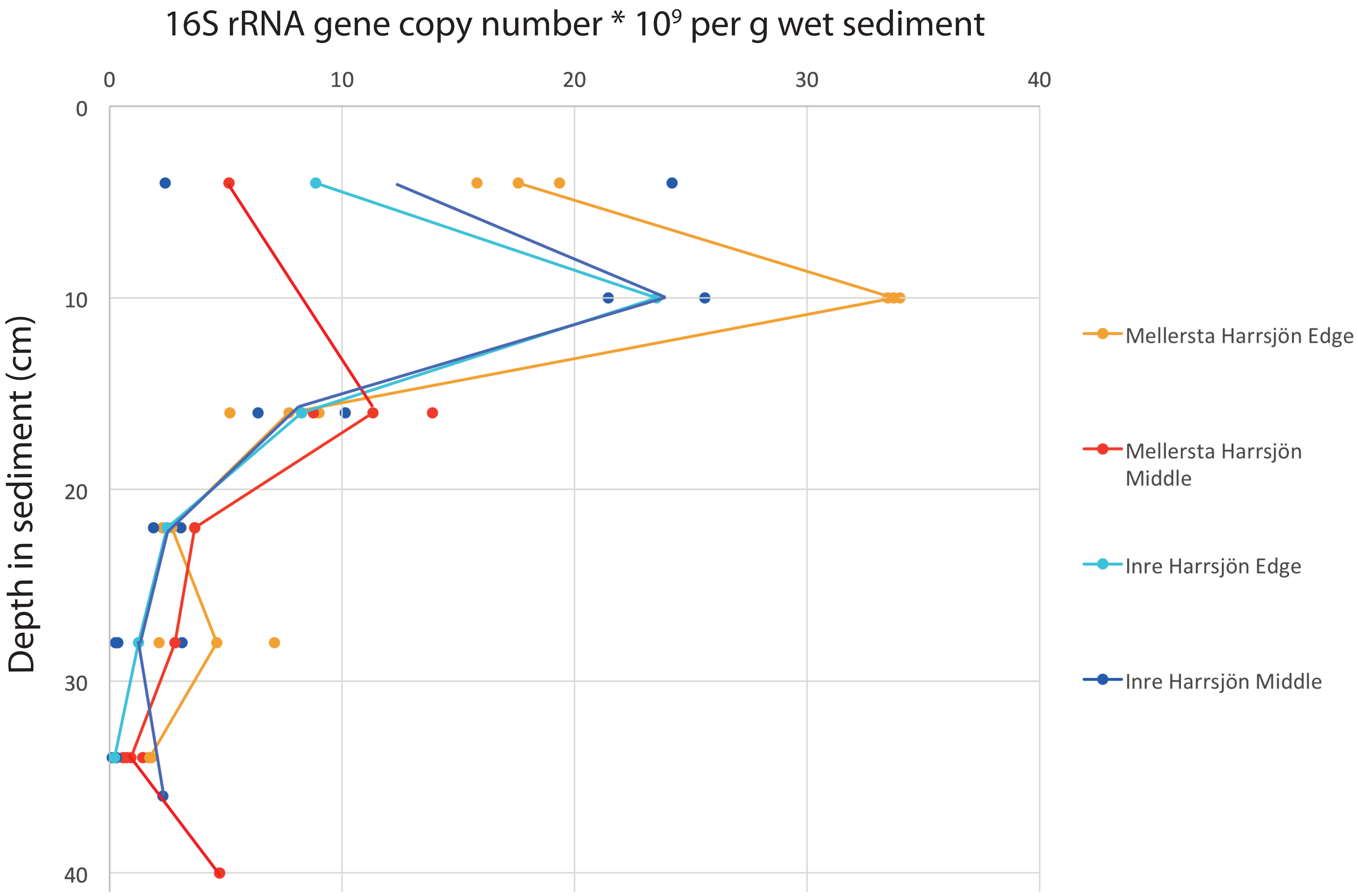


**Supplementary Figure 4. Total microbial abundances, as measured by quantitative PCR (qPCR)**. Points indicate individual measurements, lines indicate trends with depth (line placement is in the middle of replicate measurements where available). 16S rRNA gene copy number is a proxy for abundance (raw data in Supplementary Table 5).

Figure S5

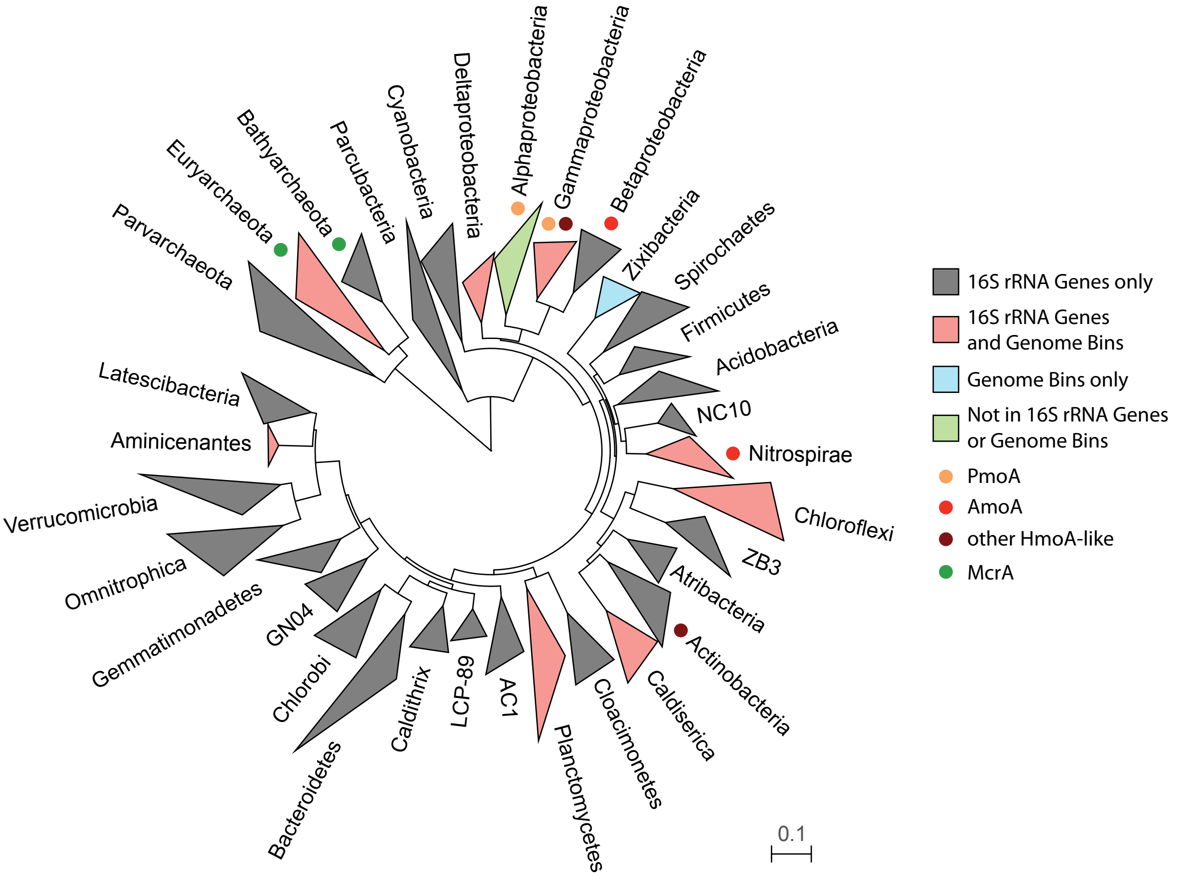


**Supplementary Figure 5. Phylogenetic tree of recovered bacterial and archaeal phyla in amplicon and metagenomic data**. Tree of representative, publicly available 16S rRNA gene sequences from lineages recovered, colored by their representation in the 16S rRNA gene amplicon data (0.5% relative abundance or higher in any sample) and/or in the 13 draft genomes reconstructed from metagenomic data (>= 50% complete, <= 10% contaminated); colored circles represent McrA (methanogenesis), PmoA (methanotrophy), AmoA (ammonia oxidation), or other HmoA-like (*e.g.*, HmoA, PxmA, function unknown but possibly related to PmoA and/or AmoA) genes recovered from metagenomic reads via the GraftM algorithm; the Alphaproteobacteria did not meet detection thresholds in the 16S rRNA gene amplicon or genomic datasets, but Alphaproteobacteria-like PmoA sequences were recovered via GraftM.

Figure S6

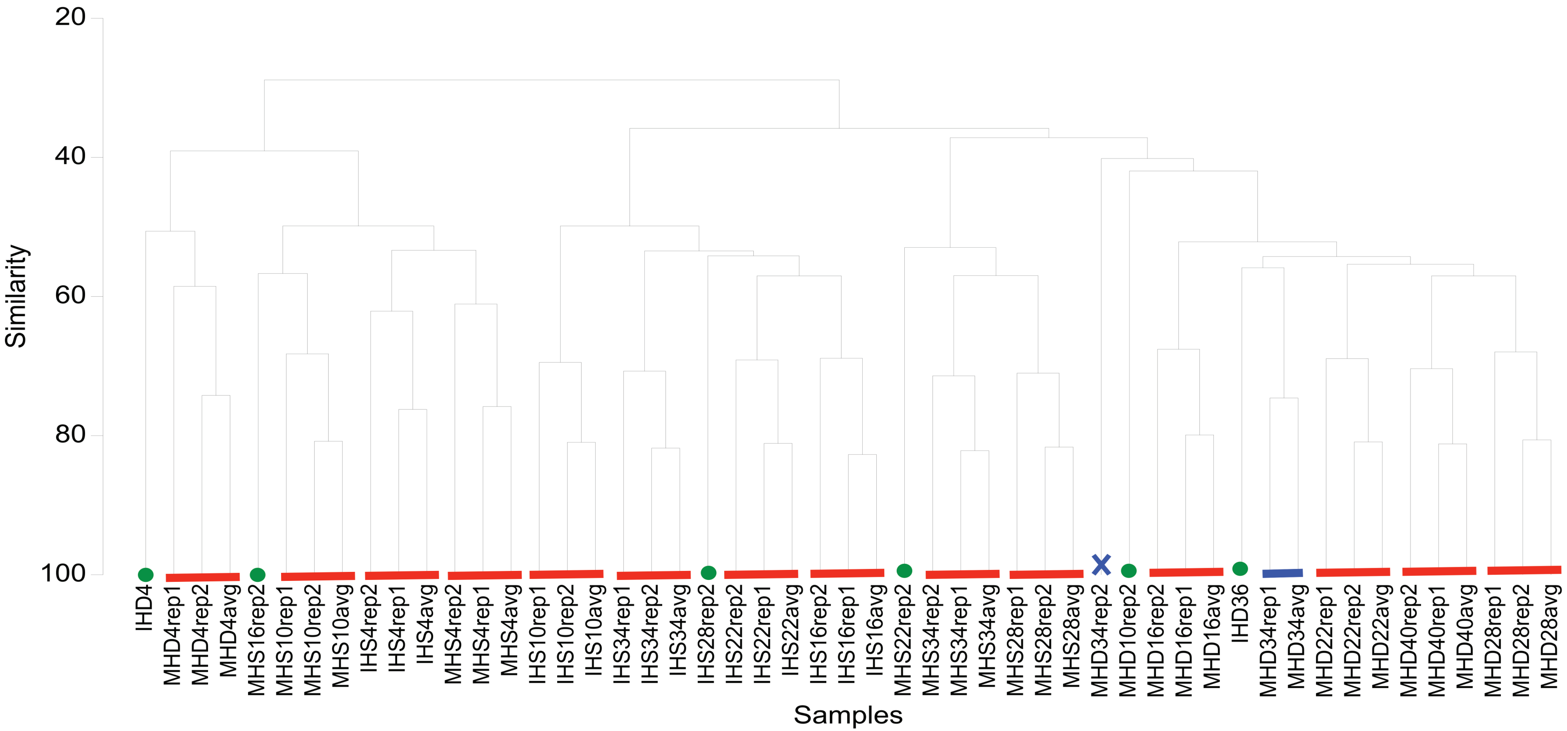


**Supplementary Figure 6.** Hierarchical clustering analysis of 16S rRNA gene-based microbial community composition in individual samples and in averages of replicates. To assess the extent to which replicates and their averages were consistent enough to use only the average values for each sample in downstream analyses, we prepared this analysis and visual representation of the underlying Bray-Curtis percent similarities (labeled “Similarity”). Red lines indicate replicate and average samples that all cluster closer to each other than to any other sample in the dataset. Green circles indicate samples with no replicates. The blue line and blue X indicate the only replicate samples that were discordant in this comparison.
